# SCSIM: Jointly simulating correlated single-cell and bulk next-generation DNA sequencing data

**DOI:** 10.1101/2020.02.03.930354

**Authors:** Collin Giguere, Harsh Vardhan Dubey, Vishal Kumar Sarsani, Hachem Saddiki, Shai He, Patrick Flaherty

## Abstract

**Background:** Recently, it has become possible to collect next-generation DNA sequencing data sets that are composed of multiple samples from multiple biological units where each of these samples may be from a single cell or bulk tissue. Yet, there does not yet exist a tool for simulating DNA sequencing data from such a nested sampling arrangement with single-cell and bulk samples so that developers of analysis methods can assess accuracy and precision.

**Results:** We have developed a tool that simulates DNA sequencing data from hierarchically grouped (correlated) samples where each sample is designated bulk or single-cell. Our tool uses a simple configuration file to define the experimental arrangement and can be integrated into software pipelines for testing of variant callers or other genomic tools.

**Conclusions:** The DNA sequencing data generated by our simulator is representative of real data and integrates seamlessly with standard downstream analysis tools.

## Background

Simulation software is important for developing and improving statistical methodology for next-generation sequencing data [1]. There are currently 149 such genetic data simulators indexed by the National Cancer Institute [2], and four of these simulators produce DNA sequencing reads with single-nucleotide variants: GemSIM [3], NEAT [4], SInC [5], and CuReSim [6]. But none of these software tools simulate bulk and single-cell data or produces data from a hierarchically grouped sampling designs typical of experimental data [7, 8].

To address this need, we have developed a software package, single-cell NGS simulator (SCSIM), to allow researchers to simulate bulk and single-cell next-generation sequencing (NGS) data from a hierarchical grouped sampling design. Figure 1 shows a high level workflow diagram of the simulator.

**Figure 1:**
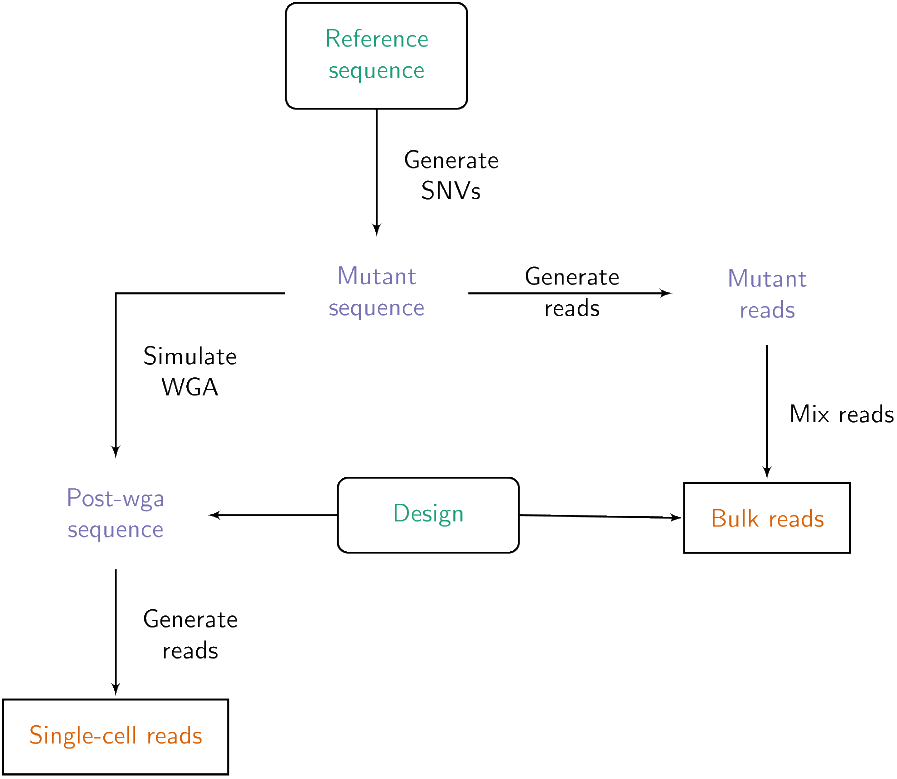
SCSIM simulation workflow. Inputs (shown in rounded boxes with green text) are the reference sequence and experiment design. Outputs (shown in cornered boxes with orange text) are the bulk and single cell FASTQ read files. Intermediate objects are shown in purple with no boxes.

The nested sampling structure is implemented using a truncated hierarchical Dirichlet process mixture model [9]. Errors induced by whole-genome amplification (WGA) of single-cells are simulated using the method described by Zafar et. al. [10]. Then, given the set of diploid reference sequences, NGS reads are simulated using `dwgsim` [11]. Finally, bulk NGS data is simulated by sampling without replacement from the set of reads from pure samples in proportions defined by the truncated Dirichlet process mixture. The command-line software takes a single haploid reference sequence in FASTA format and a YAML configuration file, and produces FASTQ reads that can be used for downstream alignment and variant calling tasks.

## Implementation

The software is implemented in python and all input parameters are controlled through a single YAML configuration file. The source code can be downloaded from the github repository https://github.com/flahertylab/scsim and the implementation can be run from a docker container defined in the repository.

### Mutated Synthetic Prototype Genome Simulation

Mutated diploid sequences are generated from a single reference (FASTA). Given *N* —the number of mutated synthetic prototype genomes and *n*—the number of possible SNV locations, one-third of SNVs are shared across all mutated sequences, one-third of SNVs are shared across one-half of the mutated sequences, and the remaining one-third of SNVs are shared across a proportion of sequences chosen from a uniform distribution [10]. The locations of the SNVs are equally spaced across the region of interest in the reference FASTA. Given the status of each SNV location in each mutated sequence, the type of diploid mutation (heterozygous or homozygous) and base substitutions are generated according to a transition probability matrix derived from Pattnaik et. al. (SiNC) [5]. The transition probabilities, and SNV locations can be set by the user, and have default values derived from literature.

### Hierarchical Sampling Structure

A hierarchical Dirichlet model is used to simulate correlation between related samples. First, the distribution over mutated synthetic prototype genomes *G*_0_ in the population is sampled from a Dirichlet with parameter *α*. Then, the distribution over mutated sequences for a biological unit *i* is sampled from a Dirichlet distribution with parameter *β*_*i*_*G*_0_ where *β*_*i*_ controls the concentration of the distribution of the biological unit around the population distribution *G*_0_. Finally, the distribution over mutated sequences for sample *j* in biological unit *i* is distributed as a Dirichlet with parameter *γ*_*ij*_*G*_*i*_ where *γ*_*ij*_ controls the concentration of the sample around the biological unit distribution *G*_*i*_.

### Single-cell Sample Simulation

For each of the *N*_sc_ single-cell samples, a whole-genome amplification (WGA) model is applied to the corresponding mutated sequence. Allelic dropout (ADO) and false positive (FP) mutations from the WGA process were generated as done previously [10]. The ADO rate was set to 20% and the FP rate was set to 3.2 *×* 10^−5^ [12]. The ADO and FP rates are calculated in reference to the entire length of the reference sequence. Finally, sequencing reads and corresponding FASTQ files were generated with `dwgsim` for each of the *N*_*sc*_ single-cell samples.

### Bulk Sample Simulation

To generate bulk sample FASTQ files where the distribution over mutated synthetic prototype genomes follows *G*_*ij*_, first `dwgsim` was used to generate FASTQ reads for each of the *n* mutated synthetic prototype genomes. Then, for each of the *N*_bu_ bulk samples, the FASTQ reads from each of the mutated synthetic prototype genomes were mixed according to the distribution *G*_*ij*_.

## Results

Here we describe simulation results showing mutations from reads generated using SCSIM are accurately identified by two different variant calling tools.

### Simulation protocol

In order to assess the accuracy and consistency of reads simulated by SCSIM, we extracted a 1 million base pair region from hg38 starting at chr20:100000. We generated three diploid synthetic prototype genomes each with 100 potential SNVs spaced every 8080 base pairs. The zygosity of the SNVs was sampled according to the method described in the implementation section. The prior parameter for the Dirichlet distribution across mutated synthetic prototype genomes, *α* was set to (0.1, 0.3, 0.6). Figure 2 shows the distribution of true SNVs across genomic position for each mutated synthetic prototype genome. The concentration parameter for the Dirichlet distribution at the biological unit level *β*_*i*_, was set to 0.1 for all *i*. The concentration parameter for the Dirichlet distribution for samples within biological units, *γ*_*ij*_ was set to 0.1 for all (*i, j*). Two samples were generated for each unit with unit 1 and 4 having one bulk and one single-cell sample, unit 2 having two single-cell samples, and unit 3 having 2 bulk samples. The mean coverage level was set to 24*×*. For bulk samples, we drew 1,000,000 reads according to the distribution over prototypes that was realized by the hierarchical Dirichlet model. A total of 90 SNVs introduced across 4 single-cell and 4 bulk samples from the described simulation protocol are the true SNVs and served as the gold standard set.

**Figure 2:**
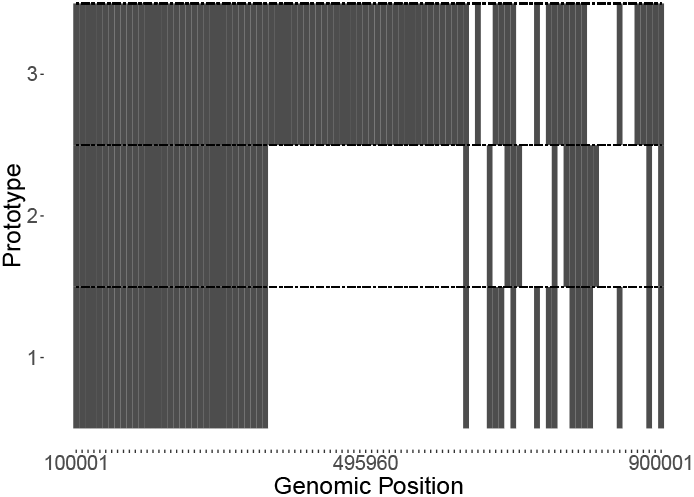
Simulated true SNV locations across mutated synthetic prototype genomes.

### Visualization of simulated reads

The simulated FASTQ reads of single-cell and bulk samples across 4 biological units were mapped to the human genome assembly (hg38) using the Burrows-Wheeler alignment tool[13]. Figure 3 shows IGV visualizations of three genomic locations with varying proportions of samples with true SNVs. Mapped reads were used to call variants by monovar[10] and BCFtools[14], two popular SNV callers used for the single-cell data. Monovar and BCFtools were run with default parameter values on the BAM files of all single-cell and bulk data. BCFtools and Monovar called 154 and 156 SNVs respectively across 4 single-cell and 4 bulk simulated samples. Our analysis showed that out of 90 true SNVs, 88 were called by both BCFtools and Monovar. Figure 4 shows a Venn diagram of the concordance between true SNVs and called SNVs from BCFtools and Monovar.

**Figure 3:**
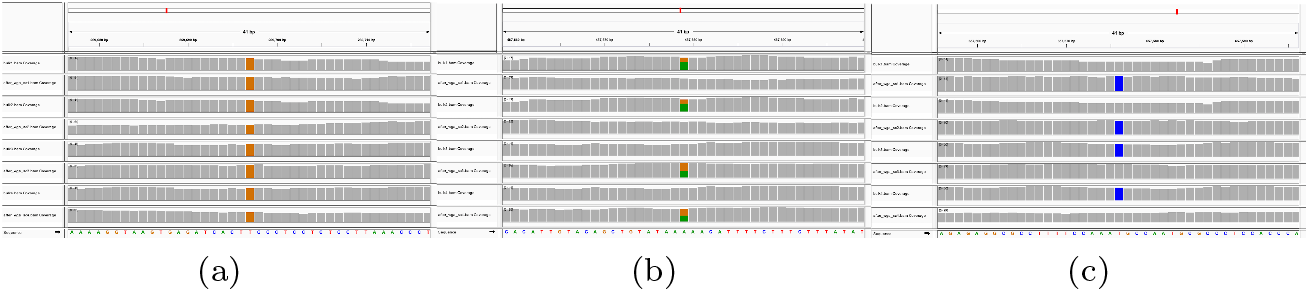
IGV visualization of reads generated by SCSIM for 4 single-cell and 4 bulk samples at (a) a genomic location where each sample has a true SNV, (b) at a genomic location where half of the samples have a true SNV, and (c) at a genomic location where a random fraction of the samples have a true SNV.

**Figure 4:**
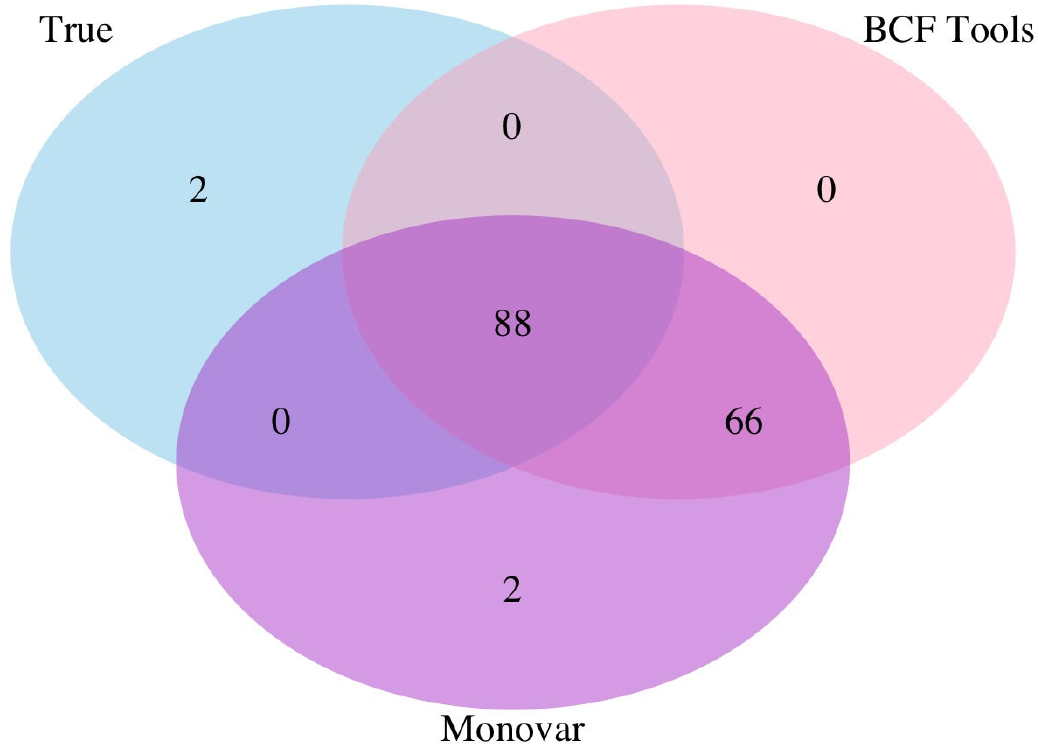
Venn diagram showing concordance of called and true variants.

## Conclusions

Given the increase in the amount of single-cell next-generation DNA sequencing data there is a need for reproducible bioinformatics methods for performing statistical inference on that data. To our knowledge there are no methods for jointly simulating bulk and single-cell sequencing data, yet these simulation tools are needed to test and validate inference methods. SCSIM jointly simulates bulk and single-cell next-generation sequencing data and generates correlated samples using a hierarchical truncated Dirichlet distribution for sampling the distribution over mutant sequences for bulk samples. Our implementation, using a docker container, allows it to be inserted in a bioinformatics pipeline without modifying existing dependencies.

## Availability and requirements

**Project name** single-cell DNA sequencing data simulator

**Project home page** https://github.com/flahertylab/scsim

**Operating systems(s)** Any

**Programming language** Python

**Other requirements** docker

**License** MIT

## Competing interests

The authors declare that they have no competing interests.

## Author’s contributions

CG contributed to the development of the algorithm and implementation and drafting of the manuscript. HVD contributed to the implementation of the algorithm, analysis and interpretation of the data, and drafting of the manuscript. VKS contributed to the analysis and interpretation of the data and drafting of the manuscript. HS contributed to the development of the algorithm. SH contributed to the implementation of the algorithm and drafting of the manuscript. PF contributed to the study conception and design, development of the algorithm, analysis and interpretation of the data and drafting of the manuscript.

## Acknowledgements

This research was funded in part by NIH award 1R01GM135931-01.

